# BCL-xL antagonizes the deleterious effects of KRAS on mitochondrial scaffolding

**DOI:** 10.1101/2022.04.13.488211

**Authors:** NM Belaid, A Basseville, G Andre-Gregoire, A Fetiveau, L Maillet, F Guillonneau, M Leduc, C Guette, L Desaubry, J. Gavard, F Gautier, PP Juin

## Abstract

In addition to its canonical role as a regulator of mitochondrial outer membrane permeabilization, BCL-xL exerts diverse non canonical functions contributing to cancer cell aggressiveness. In particular it regulates KRAS intracellular activation levels. We herein explored the mechanistic basis for this effect by a spatially restricted biotin-labelling proteomic approach designed to characterize proteins whose proximity to KRAS, used as a bait, is BCL-xL dependant. BCL-xL loss relocalizes KRAS to the vicinity of mitochondrial proteins. Proximal proteins include the mitochondrial scaffold prohibitin 2 (PHB2), which also interacts with BCL-xL and the downregulation of which prevents BCL-xL sensitive effects of KRAS induced contacts between mitochondria and endosomes, and mitochondrial mass decrease. These results argue that BCL-xL prevents a negative feedback regulation of KRAS canonical signaling by KRAS interference with mitochondrial quality control.

## Introduction

Anti-apoptotic proteins of the BCL-2 family (BCL-2, BCL-XL or MCL-1) are frequently up-regulated in cancers (Juin et al. 2013). BCL-2 homologues negatively regulate mitochondrial outer membrane permeabilization (MOMP) and resulting apoptosis by preventing activation and/or activity of multi-domain BAX/BAK downstream of BH3-only proteins, through sequestration of the BH3 domains of these pro-apoptotic counterparts. Among, BCL-2, BCL-XL and MCL-1, which exert complementary survival activities, BCL-xL binds to the widest spectrum of pro-apoptotic counterparts and this is considered to contribute, at least in part, to its potent anti-apoptotic activity. Its overexpression often correlates with chemoresistance, in particular in triple negative breast cancer (Wei et al., 2012). Due to the canonical anti-apoptotic role of BCL-xL, the presence of malignant cells with high BCL-xL expression is conceived to reflect the negative selection of low expressors by apoptotic stress. MOMP is indeed the primary way by which cancer cells die in response to radiotherapy, chemotherapy and to diverse stress stimuli cancer cells encounter as tumors progress (Juin et al. 2013). Additionally, malignant cells may also be selectively advantaged (even in the absence of pro-apoptotic pressure) as a result of non-canonical functions of BCL-xL. These functions include, non-exhaustively, the regulation of calcium signaling, of autophagy and/or, of specific transcriptional pathways. These effects were ascribed to BCL-xL ability to interact with proteins beyond the BCL-2 family (Braun et al., 2013).

We recently reported a non-canonical oncogenic effect of BCL-xL, related to its ability to favour full activation of signalling downstream of KRAS (Carné Trécesson et al., 2017). RAS pathway activation frequently occurs in solid tumors. Activation of RAS and of its downstream pathways (MAPK/ERK and PI3K/AKT) have well documented effects on cell proliferation and, in the case of epithelial cancers, on the conversion to a more migratory and invasive phenotype. We established that BCL-xL is critical for the induction by KRAS of the expression of stemness regulators and of maintenance of a cancer initiating cell (CIC) phenotype. We reported that BCL-xL interacts with KRAS, resulting in its stabilization and full activation in breast cancer cell models. The molecular mechanisms underlying this functional cooperation remain to be characterized. As BCL-xL favours the activity of constitutively GTP-bound mutant KRAS and of wild type KRAS, a regulation at the level of GTP/GDP exchange is unlikely. KRAS activity is critically determined by post translational modifications leading to the anchoring of its hydrophobic C-terminal end in subcellular membranes. While KRAS molecules are mostly localized at the plasma membrane, they are subject to intracellular trafficking events with evidence of localization on the early/late endosomal or recycling endosomal pathways (Lu et al., 2009; Cho et al., 2012). KRAS was also reported to reside at mitochondria and at contact sites between mitochondria and the ER (Bivona et al., 2006; Sung et al., 2013). It is currently unclear how exactly this compartmentalization influences the duration and output of canonical and non-canonical KRAS signals.

To unravel which elements of the complex regulation of KRAS signaling are influenced by BCL-xL we sought to characterize changes in the intracellular context of the former upon modulation of the latter. We reasoned that assumption-free mapping of endogenous proteins in the vicinity of KRAS in a cellular context expressing BCL-xL or not would be highly informative in this context. We thus developed a site restricted enzymatic tagging approach using KRAS as a bait introduced in CRISPR edited breast cancer cells to identify KRAS proximal proteins by quantitative proteomic analysis. Our results put forth that the cooperation between KRAS and BCL-xL stems from the latter’s ability to antagonize a previously unforeseen tumor suppressive effect of the former, related to interference with mitochondrial scaffolding.

## RESULTS

### An unbiased proteomic approach reveals a major impact of BCL-xL on KRAS proximitome

BCL-xL influence on KRAS signaling may stem from the ability of the former to influence elements of the intracellular context which are critically required for the latter to sustain downstream signaling (and maintain protein expression levels). To gain insight into these, we used a KRAS construct fused at its N-terminal end with APEX, an ascorbate peroxidase derivative relying on hydrogen peroxide for catalyzing the oxidation of biotin-phenol to a short-lived and reactive biotin-phenol free radical which reacts with electron-rich amino acids on neighboring proteins resulting in their biotinylation (Lee et al., 2016). KRAS proximitomes were identified by quantitative mass spectrometric analysis of endogenous proteins specifically present in streptavidin pull downs following transient transfection of the APEX-KRAS bait in MCF-7 cells, APEX enzymatic activation by treatment of resulting cells with H202 (1 min), in the presence of biotin-phenol, prior to cell lysis (Figure 1A). We used MCF-7 cells as these cells showed optimal transfection under the conditions used (80% of cells positive for YFP fluorescence following transfection with YFP-KRAS), and since we established that BCL-xL interferes with KRAS signaling in this context (Carné Trécesson et al., 2017). To investigate the influence of BCL-xL on KRAS proximitome, we performed experiments using MCF-7 where BCl-xL was deleted by CRISPR/CAS9, together with control cells. As shown in Supplementary Figure 1, BCL-xL deleted MCF-7 showed enhanced sensitivity to cytochrome c release induction (as judged by flow cytometry) by treatment with the MCL-1 antagonist S63845. This indicates that endogenous BCL-xL exerts in these cells its canonical function to preserve MOMP together with its complementary homolog MCL-1. Further validating our deletion strategy, BCL-xL deleted cells proved less able to transmit signals from transiently transfected YFP-KRAS (as judged by phosphorylation of ERK) at baseline or in response to EGF, even when comparable amounts of YFP-KRAS were obtained by the transduction procedure (Figure 1B).

**Figure 1.**
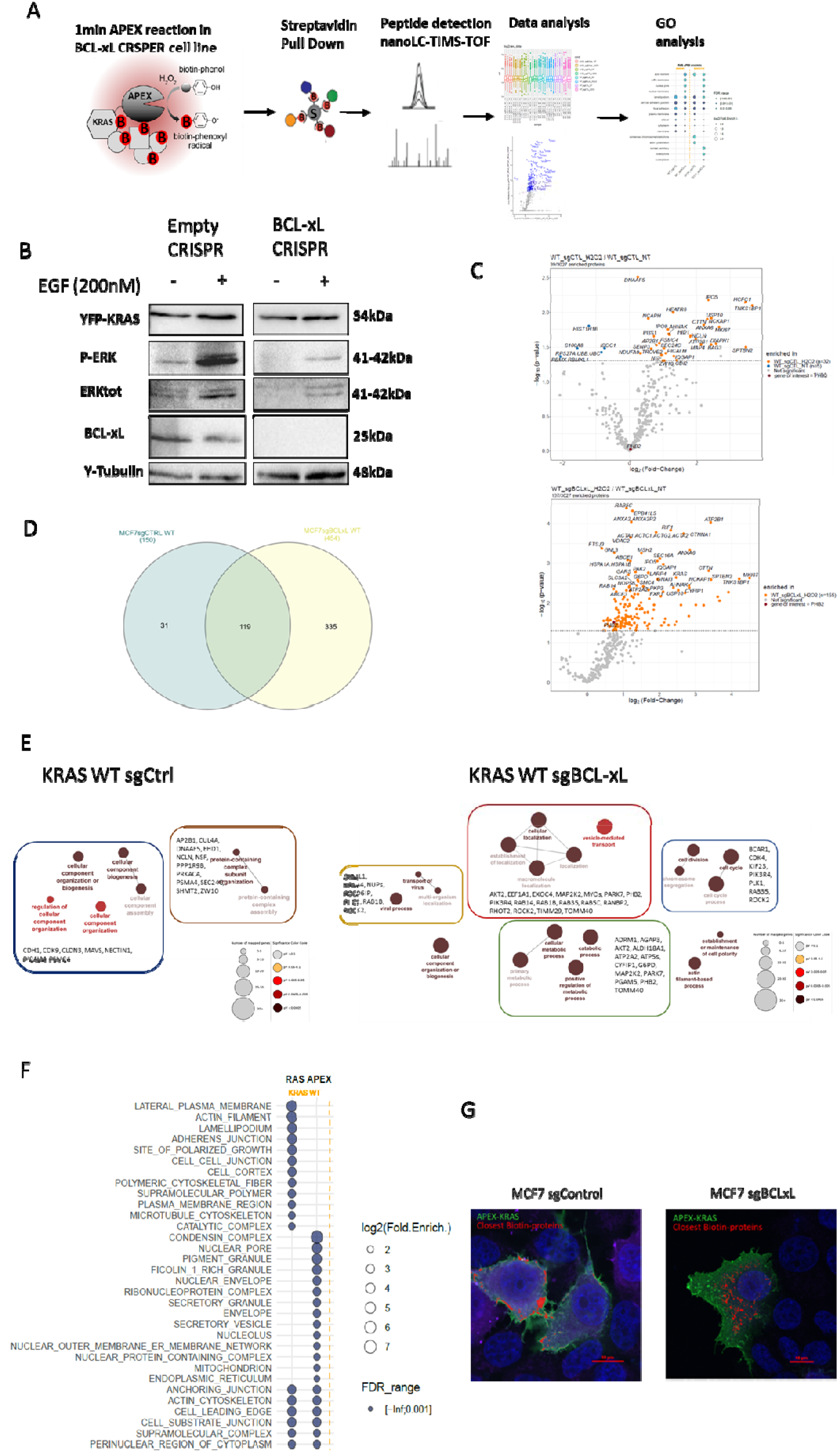
A unbiased proteomic approach reveals a major impact of BCL-xL on KRAS proximitome. **A** Representation of APEX protocol analysis. APEX biotinylation reaction is induced by H2O2 addition in cell medium in MCF7 Empty CRISPR or in MCF7 BCL-xL CRISPR cell lines. To exclude non-specific partners, H2O was added instead of H2O2 as a negative control. Biotinylated proteins were purified using streptavidin-coupled magnetic beads and detected by mass spectrometry (nanoLC-TIMS-TOF). **B** Western Blot analysis of KRAS activity in MCF7 Empty CRISPR or in MCF7 BCL-xL CRISPR cell lines expressing YFP-KRAS. Representative of three independent experiments. **C** Vulcano Plot visualization of selected proximitome in MCF7 Empty CRISPR or in MCF7 BCL-xL CRISPR cell lines. **D** Venn Diagramm visualization of selected protein lists. **E** ClueGo representation of Gene Ontology biological processes. For each cell line, only non-common partners (shown in D) were analyzed using GO enrichment analysis. **F** Comparative analysis of KRAS localization prediction. For each indicated context, subcellular enrichment analysis with GSEA were performed on each specific protein list. **G** Confocal microscopy in MCF7 Empty CRISPR or in MCF7 BCL-xL CRISPR cell lines after APEX-KRAS biotinylation reaction. APEX-KRAS in labeled in green through anti-Flag antibody, Biotinylated Proteins in DeepRed using Streptavidin coupled to Alexa647 and a colocalization analysis was performed using Nikon Software, represented in Red. Representative of three independent experiments.

To characterize the BCL-xL dependent KRAS proximitome, we performed four independent transient transfection assays in two distinct cell contexts (BCL-xL proficient or deficient). Resulting cells were treated with H_2_O_2_ for one minute or left untreated to provide for each context four pairs of tests with their matched negative controls. The presence of a bit more that 3 400 proteins was quantified in each of the 16 assays. Proteins were dubbed as being specifically part of the KRAS proximitome when following one of these three criteria:

i. if their presence was detected in four positive tests compared to none or only one in the negative controls
ii. if their presence was detected in three positive tests compared to none in the negative control
iii. if they were quantified at significantly higher levels in the tests compared to the negative control by Vulcano plot analysis (Figure 1C)

In a last step, we deleted from the two KRAS proximitome lists protein members of the “crapome” (Mellacheruvu et al., 2013), as promiscuous ability to be pulled down by numerous baits casts doubt on the identification as a *bona fide* KRAS proximal protein by our approach.

As a result, we identified 150 proteins specifically present in the vicinity of KRAS in control MCF-7 cells and 454 ones in BCL-xL deleted cells (Figure 1C). While 119 proteins were commonly found in each condition, 31 were specific for the control condition and 335 were specifically found in BCL-xL deficient cells (Figure 1D) indicating a major qualitative change in the proteomic context of KRAS when BCL-xL is absent. Gene ontology analysis indicated that proteins specifically found in the vicinity of the KRAS bait in BCL-xL deleted cells are enriched for proteins involved in the following processes: cell cycle, catabolism, transport of virus and cellular localization. Moreover, bait localization prediction by Gene Set enrichment analysis hinted on a preferential localization at the plasma membrane of the KRAS proximitome in control cells (as indicated by biotin labelling of E-cadherin, CDH1), and at nuclear membranes, early endosomes and mitochondria in BCL-xL deleted cells.

These qualitative modifications indicate that steady state localization of KRAS is modified upon BCL-xL loss. To test this, we used confocal microscopy to perform by immunofluorescence colocalization analysis of APEX-KRAS and proteins biotinylated following APEX activation. We reasoned that the localization of proteins which are closest to APEX-KRAS would reflect where, within the cell, bait molecules spend most of their time. As shown in Figure 1F, whereas this approach revealed a stable plasma membrane localization of APEX-KRAS in control cells, it revealed an intracellular localization for APEX-KRAS in BCL-xL deficient cells.

### Upon BCL-xL deletion, KRAS accumulates to mitochondria in the vicinity of PHB2

We focused on the enhanced mitochondrial localization of KRAS inferred by our proteomic approach. Indeed, 51 out of the 454 proteins characterized in the vicinity of KRAS in BCL-xL deleted cells are in the list of 1 555 proteins referenced as possible mitochondria residing ones (Morgenstern et al., 2017) as opposed to 12 proteins (out of 150) in control cells (Figure 2A left). This corresponds to a net enrichment in the proportion of mitochondrial proteins in the proximitome of KRAS upon BCL-xL loss (Figure 2A, right). Enhanced localization of transiently transfected YFP-KRAS in BCL-xL deleted cells to mitochondria was consistent with its accumulation in heavy membrane fractions (where BCL-xL localizes in control cells, and marked by COX IV expression) obtained by subcellular fractionation assays (Figure 2B). These assays revealed that endogenous KRAS also accumulates at heavy membrane fractions in the absence of BCL-xL. To further confirm that KRAS exhibits an enhanced mitochondrial localization upon BCL-xL loss, we performed confocal microscopic analysis of transiently transfected cells and measured colocalization indexes between YFP-KRAS and mitotracker staining. Distance map analysis using Saito’s method (Saito and Toriwaki, 1994) was performed to calculate two channels-pixels proximity. This method revealed that KRAS was closer to mitochondria in BCL-xL deficient cells than in BCL-xL proficient cells (Figure 2C).

**Figure 2.**
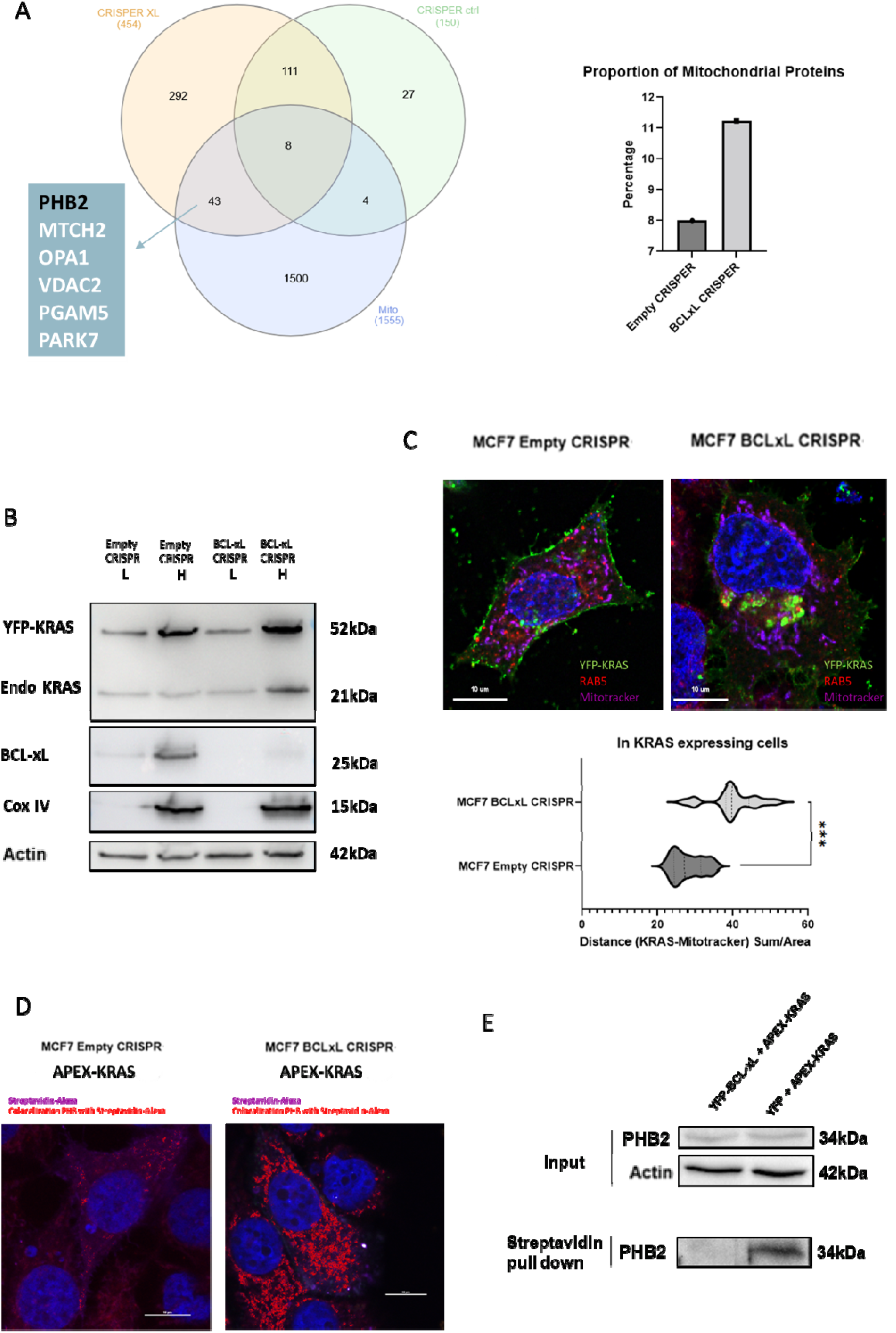
Upon BCL-xL deletion, KRAS accumulates to mitochondria in the vicinity of PHB2. **A** Venn Diagram of KRAS proximitomes in both Emtpy CRISPR and BCL-xL CRISPR cell lines with known mitochondrial proteins (Left). Proportion of mitochondrial proteins in each list is represented as a histogram (Right). **B** Western Blot analysis of Light (L) or Heavy (H) fractions after digitonin extraction in MCF7 Empty CRISPR or in MCF7 BCL-xL CRISPR cell lines expressing YFP-KRAS. Representative of three independent experiments. **C** Confocal microscopy in MCF7 Empty CRISPR or in MCF7 BCL-xL CRISPR cell lines expressing YFP-KRAS. YFP-KRAS (green), Rab5 (Red) and Mitotracker (DeepRed). Distance Map analysis of KRAS distance from MitoTracker. Representative of three independent experiments. **D** Confocal Microscopy in MCF7 Empty CRISPR and MCF7 BCL-xL CRISPR cell lines after APEX-KRAS biotinylation reaction. Biotinylated proteins (DeepRed), Colocalization of Biotinylated proteins and PHB2 (Red) using Nikon software. Representative of three independent experiments. **E** Western Blot analysis of lysates (Input) and biotinylated proteins (Streptavidin pull down) after APEX-KRAS transfection/activation in cells over-expressing YFP-BCL-xL or YFP alone. Representative of three independent experiments. ***: p-value <0,0001

String database analysis of the 43 mitochondrial proteins that were specifically detected in the proximity of KRAS in BCL-xL deficient cells (Figure 2A left) revealed a group of proteins known to interact with each other and involved in mitochondrial scaffolding and function, with a critical contribution of PHB2 (Supplementary Figure 2). Confocal microscopy analysis of the colocalization of biotinylated proteins (following APEX-KRAS transfection/activation) and of endogenous prohibitins was performed. This revealed enhanced vicinity between KRAS proximal proteins and prohibitins (potentially at subcellular membranes) upon BCL-xL loss (Figure 2D). This indicates that under these conditions the steady state localization of the KRAS bait is more frequently in the proximity of prohibitins. PHB2 seemed to us of particular interest in this study as we also specifically identified it following mass spectrometric analysis of YFP-BCL-xL immunoprecipitates from MCF-7 cells stably expressing this fusion protein (not shown), together with ATP5C1, ATP5L and VDAC2 (which were reported to interact with PHB2 and which are, akin to the latter, in the list of 43 KRAS proximal mitochondrial proteins identified in BCL-xL deficient cells). Candidate site restricted enzymatic tagging assays, based on western blot analysis of PHB2 in streptavidin pull downs from APEX-KRAS transfected YFP-BCL-xL or YFP expressing MCF-7 cells revealed that overexpressed YFP-BCL-xL prevents KRAS-PHB2 proximity. (Figure 2E).

### BCL-x loss enhances the presence of KRAS at the interface between mitochondria and endosomes in a PHB2-dependent manner

Our detection of an enhanced proximity between RAB5 and KRAS upon BCL-xL loss in proteomic analysis led us to investigate KRAS contacts with endosomes further. Microscopic analysis revealed an increase of YFP-KRAS colocalization with RAB5 upon BCL-xL loss, as judged by an enhanced Pearson coefficient between YFP and RAB stainings in BCL-xL deleted cell populations. In sharp contrast, the distance between YFP-KRAS and the recycling endosome marker Rab11 was unaffected by BCL-xL loss (Figure 3A).

**Figure 3.**
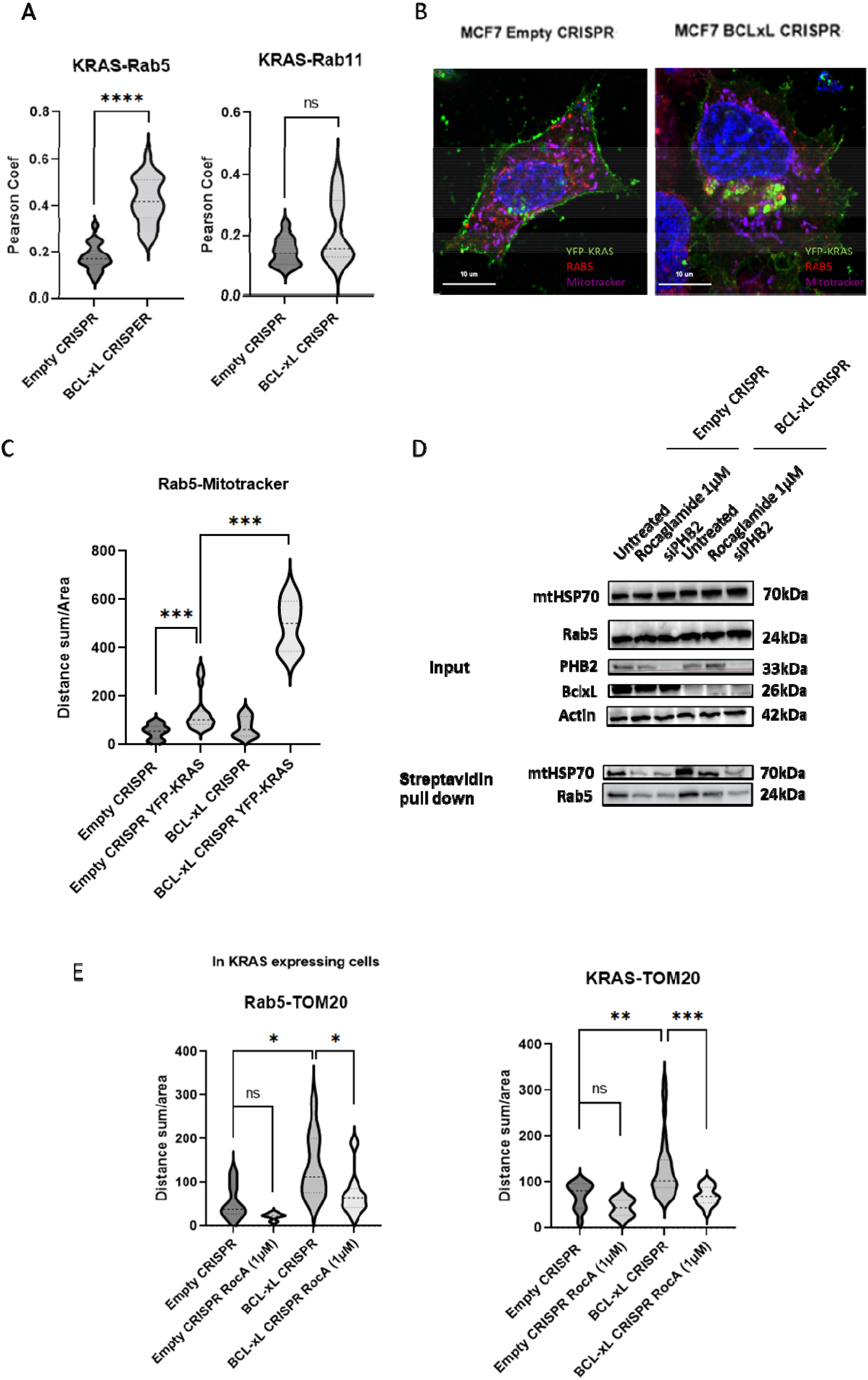
BCL-x loss enhances the presence of KRAS at the interface between mitochondria and endosomes in a PHB2 dependent manner. **A** Colocalization analysis of KRAS with the early endosomal marker Rab5 and with the recycling endosome marker Rab11. Pearson Coefficient was obtained using ICY software. N=200 cells per condition. **B** Confocal microscopy in MCF7 Empty CRISPR or in MCF7 BCL-xL CRISPR cell lines expressing YFP-KRAS. YFP-KRAS (green), Rab5 (Red) and Mitotracker (DeepRed) are shown. Representative of three independent experiments. **C**. Distance Map analysis of Rab5 distance from Mitotracker in MCF7 Empty CRISPR or in MCF7 BCL-xL CRISPR cell lines expressing or not YFP-KRAS. N=200 cells per condition. **D** Western Blot analysis of biotinylated proteins after APEX-KRAS biotinylation in MCF7 Empty CRISPR and MCF7 BCL-xL CRISPR and Streptavidin pull down. Representative of three independent experiments. **E** Distance Map analysis of Rab5 and KRAS distance from the mitochondrial marker TOM20 in MCF7 Empty CRISPR and MCF7 BCL-xL CRISPR expressing YFP-KRAS and treated of not with 1μM Rocaglamide for 18h. N=50 cells per condition. ns: P value>0,05 ***: P value <0,0001 ****: P value <0,0001

We inferred that the coincidence of KRAS localization at mitochondria and at endosomes in BCL-xL deleted cells might be due to enhanced contact between the two organelles as seen by triple staining for YFP-KRAS, Rab5 and Mitotracker (Figure 3B). Thorough measurement of RAB5-Mitotracker contacts in cell populations indicated that they significantly increased upon YFP-KRAS expression in the absence of BCL-xL (Figure 3C).

We explored a role for PHB2 in KRAS contacts with mitochondria and endosomes. To do this, we downregulated its expression by siRNA in control and BCL-xL deficient cells, prior to a proximity labelling assay based on the transient transfection of APEX-KRAS. Western blot analysis in streptavidin pull downs for mtHSP70 expression revealed that KRAS proximity with this mitochondrial marker, which was enhanced in BCL-xL deleted cells, was decreased by PHB2 downregulation (Figure 3D). PHB2 downregulation also decreased the BCL-xL deletion sensitive proximity between KRAS and RAB5 (Figure 3D). Pharmacological modulation of prohibitins using Rocaglamide (Polier et al., 2012) recapitulated the effects of siPHB2 on KRAS presence in the vicinity of mtHSP70 and Rab5 (Figure 3D). Preliminary investigation, by distance mapping analysis of confocal microscopic images, of transiently transfected KRAS, endogenous RAB5 and TOM20 localizations confirmed that rocaglamide treatment antagonized the enhanced mitochondria to endosomes contacts (Figure 3E) as well as KRAS at mitochondria (Figure 3E). Taken together, these observations argue that KRAS interactions with PHB2 contribute to its localization to mitochondria, and that the latter is necessary for its recruitment in the vicinity of endosomal RAB5.

### KRAS promotes PHB2 dependent mitochondrial clearance and increases mitovesicle production

Mitochondrial prohibitin complexes contribute to mitochondrial quality control. Morevoer, a Rab5 endosomal pathway is involved in sensing mitochondrial stress (Hammerling et al., 2017). We thus hypothesized that the aforementioned observations result from some mitochondrial damage induced by KRAS. Evaluation of TOM20 staining intensity (as a readout of mitochondrial mass) indicated that YFP-KRAS expression in control cells tended to lead to the appearance of a cells with low TOM20 levels and that YFP-KRAS expression in BCL-xL deficient cells had a major inhibitory effect on TOM20 levels (Figure 4A). FACS analysis of TOM20 expression under the same conditions led to a similar conclusion (Figure 4B). This argues that KRAS promotes BCL-xL sensitive mitochondrial clearance.

**Figure 4.**
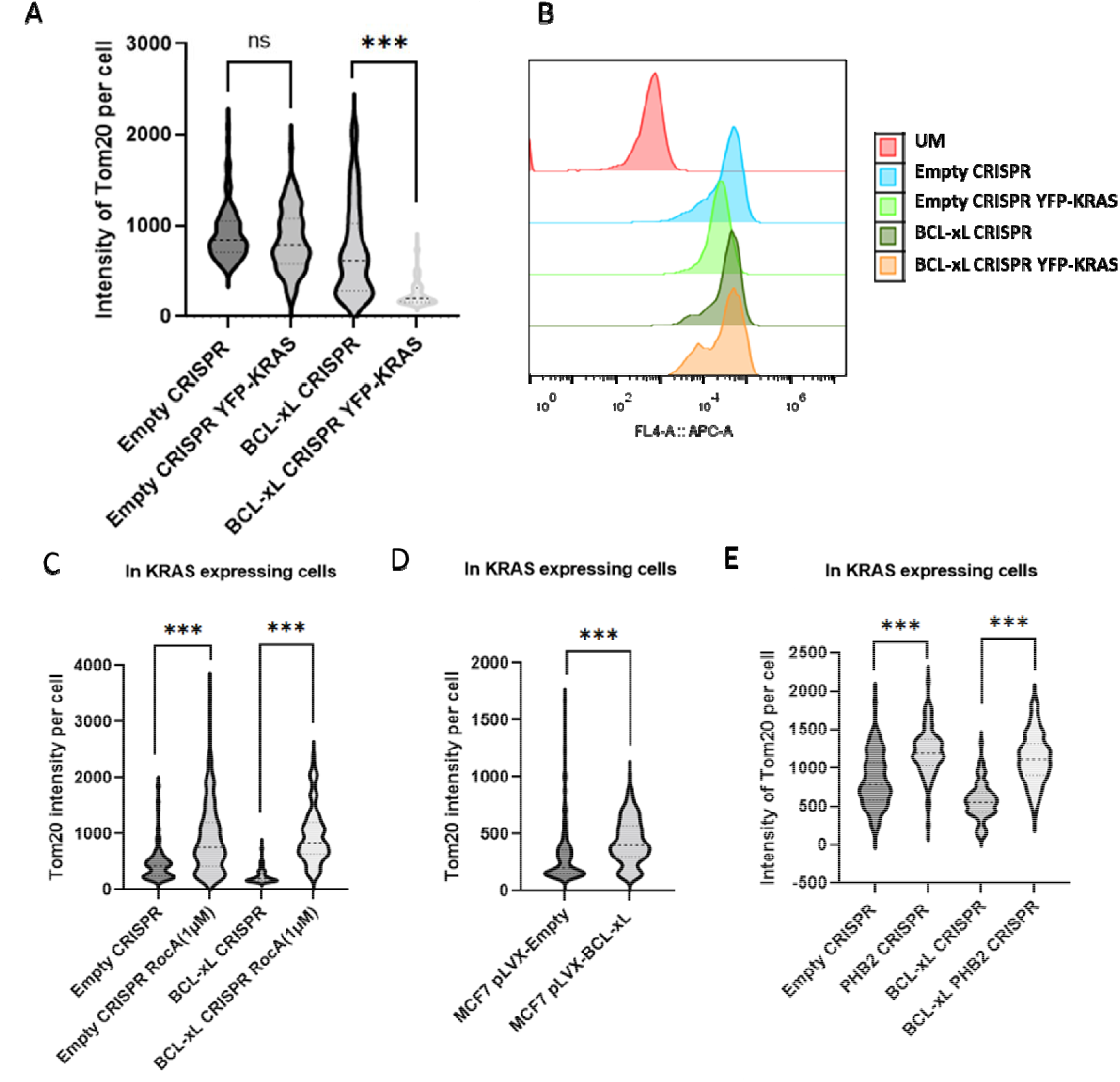
KRAS promotes PHB2 dependent mitochondrial clearance and increases mitovesicle production. **A** Confocal image analysis of TOM20 intensity per cell, obtained using ICY software, in MCF7 Empty CRISPR or in MCF7 BCL-xL CRISPR cell lines expressing or not YFP-KRAS. N=200 cells per condition. **B** Mitochondrial Mass estimated by TOM20 fluorescent intensity measured by Flux Cytometry. Representative of three independent experiments. **C**. Confocal image analysis of TOM20 intensity per cell, obtained using ICY software, in MCF7 Empty CRISPR or in MCF7 BCL-xL CRISPR cell lines expressing YFP-KRAS and treated or not 18h with 1μM of Rocaglamide. N=100 cells per condition. **D** Confocal image analysis of TOM20 intensity per cell, obtained using ICY software, in MCF7 stably expressing pLVX-Empty or pLVX-BCL-xL cell lines expressing YFP-KRAS. N=200 cells per experiment. **E** Confocal image analysis of TOM20 intensity per cell, obtained using ICY software, in MCF7 Empty CRISPR, MCF7 PHB2 CRISPR, MCF7 BCL-xL CRISPR or MCF7 double Knock Out BCL-xL and PHB2 CRISPR cell lines expressing YFP-KRAS. N=200 cells per condition. ns: P value>0,05 ***: P value <0,0001

To investigate a role for prohibitins in KRAS induced loss of mitochondrial mass, we investigated TOM20 staining intensity in YFP-KRAS expressing BCL-xL proficient and deficient cells treated or not with rocaglamide. This prohibitin ligand induced TOM20 expression in BCL-xL deficient cells (Figure 4C). Interestingly, it also induced mitochondrial mass increase in KRAS expressing control cells, indicating that mitochondrial clearance is antagonized by BCL-xL, but that it can occur even in its presence. Consistent with this, BCL-xL overexpressing cells maintained higher mitochondrial (TOM20) levels than control cells upon expression of YFP-KRAS (Figure 4D). To address a role for PHB2 by a genetic approach with confocal microscopy, we attempted to delete the PHB2 gene in control and BCL-xL deleted cells by CRISPR/CAS9 (to ensure that each analyzed cell is deleted). This deletion proved lethal to these cells in the long term and we thus could only investigate cells at early stages, setting aside for the analysis these with overt morphological damage. Consistent with the rocaglamide treatment results, investigation of healthy PHB2 deleted cells revealed their resistance to KRAS induced TOM20 decrease in the absence of BCL-xL (Figure 4E).

## Discussion

In this work our investigation of the consequence of BCL-xL depletion on KRAS proximitome led us to unravel an influence of KRAS on mitochondria, with which it can directly interact. Preceding reports have established that KRAS4b can localize at this organelle upon phosphorylation of S181 within its polybasic region. This leads to a mitochondrial interaction with BCL-xL that promotes mitochondrial outer membrane permeabilization (Bivona et al., 2006). This favors also an interaction, presumably at contact sites between mitochondria and ER, with inositol triphosphate receptors (Sung et al., 2013). Even if we detected biotinylation of ITPR3 following expression of APEX-KRAS in BCL-xL depleted MCF-7 cells, (data not shown), it should be noted that we used KRAS4a, which does not harbor a phosphorylable S181 residue. In our study, moreover, KRAS mitochondrial localization occurs in the absence of BCL-xL, and not through a direct interaction with this protein, suggesting that the mechanism is different. The consequence of KRAS localization at the mitochondria in the absence of BCL-xL is also different from that reported in preceding studies as we detected a decrease in mitochondrial mass (as judged by TOM20 expression). KRAS-4a was already described to interact in a depalmitoylation dependent manner with mitochondrial hexokinases and with VDAC (Amendola et al., 2019). Whether hexokinases are involved, and whether the mitochondrial localization of KRAS in our experiments coincides with a metabolic effect, remain to be characterized.

Mitochondrial proteins identified in the vicinity of KRAS upon BCL-xL-depletion present different sub-localizations, including outer membrane (TOMM40, MTCH2, VDAC2), inner membrane (PHB2, STOML2) and matrix (mtHSP70). This suggests profound rearrangements in the architecture of mitochondria under these conditions and is consistent with the eventual mitochondrial mass decrease observed. Mitochondrial damage under these conditions would explain induced proximity of KRAS and Rab5 at contacts sites between mitochondria and endosome. Indeed, a Rab5 endosomal pathway is involved in sensing mitochondrial stress and promoting an adaptative response (Hammerling et al., 2017). Other BCL-xL dependent KRAS proximal proteins are known actors of mitochondrial quality control. PARK7 is a direct effector of autophagy (Lee et al., 2018) while PHB2 was established to bind to the autophagosomal membrane-associated protein LC3 (MAPLC3) through an LC3-interaction region (LIR) domain and to play an essential role in Parkin dependent mitophagy by a PGAM5 dependent process (Yan et al., 2020).

While our data strongly advocate that BCL-xL prevents KRAS from directly inducing mitochondrial dysfunction, linked to mitochondrial clearance, the exact mechanisms accounting for mitochondrial mass decrease upon KRAS expression remain to be determined. We used a model cell line that notoriously exhibits low PARKIN expression (Liu et al., 2017). However, a process similar to that through which mitochondrial Aurora Kinase A triggers mitophagy in a PARKIN independent yet PHB2/MAP1LC3B dependent manner might be involved (Bertolin et al., 2021). Our preliminary data do not support this hypothesis. Using the mitoTandem probe in BCL-xL depleted MCF-7 transiently transfected with KRAS (data not shown), we were unable to detect inclusion of mitochondria in lysosomes, which is a necessary final step for mitochondrial degradation by (PARKIN dependent or independent) mitophagy or by the Rab5 endosomal pathway. We were also unable to detect MAP1LC3 in the proximity of KRAS in any of the conditions used. It should be noted that induction of autophagy by phosphorylated KRAS was found to be lost in BCL-xL knock out mouse embryonic fibroblasts (Sung et al., 2013). This is somehow consistent with the reported pro-autophagic role for BCL-xL in KRAS mutant colorectal cancer cells (Priault et al., 2010) and suggests that cells used in our study may not be fully equipped to target mitochondrial components to lysosomes. They are nevertheless able to promote mitochondrial clearance in response to mitochondrial dysfunction, as indicated by the decrease in TOM20 levels we could detect by flow cytometry upon mitochondrial potential collapse induced by CCCP (not shown).

A stable relocalization of KRAS to intracellular membranes upon BCL-xL loss may directly contribute to decrease its canonical signaling output. In the context of EGFR stimulation in particular, activation of downstream effectors critically occurs, indeed, at the plasma membrane (Surve et al., 2019). It is also plausible that the decrease in RAS activation is also fueled by a destabilization of intracellular KRAS expression. Consistent with this, we found by cytometry that YFP-KRAS tends to be expressed at lower levels in BCL-xL depleted cells (not shown). Our study puts forth the following hypothesis to account for this effect, which we already reported (Carné Trécesson et al., 2017): BCL-xL may not regulate baseline KRAS turnover, but BCL-xL would instead mitigate deleterious effects of KRAS on mitochondria, that would lead to a decrease of mitochondrial proteins together with KRAS itself. This would occur through mitochondrial damage induced degradation processes that may differ depending on cellular contexts, and that require further characterization.

PHB2, which is found in the vicinity of KRAS, and which tightly colocalizes to APEX-KRAS biotinylated proteins upon BCL-xL loss seems to play a central role in mitochondrial damage unraveled by our study. Indeed, its downregulation prevents proximity between KRAS and mtHSP70, with Rab5 and concomitantly prevents mitochondrial mass decrease. Mitochondrial prohibitins PHB and PHB2 belong to the stomatin/prohibitin/flotillin/HflKC (SPFH) family of membrane anchored proteins. In addition to the role for PHB2 as an actor of mitophagy mentioned above, PHB/PHB2 heterodimers play a key role in mitochondrial quality control through regulating mtDNA maintenance, protein synthesis and degradation, assembly of the OXPHOS complexes and maintenance of cristae structures (reviewed in Hernando-Rodríguez et Artal-Sanz 2018). PHB complexes have a rich interactome of intermembrane space proteins contributing to mitochondrial scaffolding and dynamics, and complex assembly critically relies on a coiled-coil region in PHB2 (Yoshinaka et al., 2019). KRAS may interfere with these complexes to promote mitochondrial dysfunction. We think that this occurs regardless of the GTP binding state of KRAS. Indeed, assumption free investigation of APEX-KRASV12 proximitome in BCL-xL proficient or deficient MCF-7 cells also revealed enhanced proximity between KRASV12 and PHB2 upon BCL-xL deletion (note that we observed enhanced proximity with OPA1, a known interactant of PHB2). Moreover, preliminary flow cytometric analysis indicates that transiently transfected KRASV12 induces a decrease in TOM20 expression and that it does so all the more as BCL-xL is depleted (not shown).

Even though the molecular basis for KRAS recruitment to mitochondrial PHB complexes remains elusive, it may be similar to that involved in its recruitment to PHB complexes at the plasma membrane, that positively regulate downstream oncogenic signaling (Rajalingam et al., 2005). This similarity is suggested by the fact that rocaglamide, which interferes with plasma membrane PHB (Yurugi et al., 2017) also prevents KRAS proximity with mtHSP70 and Rab5 in the absence of BCL-xL. This effect recalls the inhibitory effect of another PHB ligand, xanthohumol, on Aurora Kinase A induced mitochondrial degradation (Bertolin et al., 2021). This underscores that modulation of RAS activity by prohibitin targeting may have ambiguous effects: inhibition of canonical RAS signaling may be balanced by the inhibitory effects on RAS induced mitochondrial dysfunction. As BCL-xL expression limits the latter, one may assume that BCL-xL will critically determine the net biological effect of prohibitin targeting.

A general implication of our study is that malignant cells rely, to fully support oncogenic RAS signaling, on robust mitochondrial homeostatic programs. How BCL-xL participates to such programs remains to be determined, as neither MOMP nor autophagy seem to be patently triggered under our conditions. In MCF-7 cells possible interactants of BCL-xL (data not shown) include pro-apoptotic proteins of the BCL-2 family and a group of outer membrane (VDAC2) and inner membrane proteins (ATP5 subunits, PHB2) reminiscent of PHB2 containing mitochondrial complexes (Yoshinaka et al., 2019) in addition to PHB2 itself. BCL-xL might participate to such complexes in a similar manner to MAVS, a protein involved in inflammatory signaling with a comparable outer membrane anchoring by a C-terminal end, and somehow contribute to their stability and maintenance function. The presence of BCL-xL in these mitochondrial scaffolds also implies that the latter might reciprocally influence the canonical function of the former. In agreement with this, targeting mitochondrial structure was shown to enhance sensitization to antagonists of the BCL-xL homolog BCL-2 (Chen et al., 2019).

## Methods and Material

### Cell culture and reagents

MCF7 cells lines were obtained from ATCC and modified through viral infection of pLentiCRISPR V2 plasmid (with or without a guide sequence) and puromycin or hygromycin selection. MCF-7 stably overexpressing YFP-BCL-xL or YFP were obtained by transfection with peYFP-BCL-xL and selection with G418. Cells were cultured in RPMI 1640 (Gibco, Fisherscientific) supplemented in 10% SVF (Eurobio), 1% Glutamin and 1% Penicillin/Streptomycin. Cells were grown at 37C° in 5% CO_2_. For several experiments, including siRNA experiments, transfections were performed using Lipofectamine 2000 according to the manufacturer’s instructions. For siRNA experiments, ON-TARGETplus Human PHB2 siRNA was purchased from Dharmacon and 5nM was transfected per condition in 6-well plates. Cells were treated with 1μM of Rocaglamide (HY-19356, MedChemExpress) in cell medium.

CRISPR Guide sequences:

BCL-xL: 5′-CACCGGCAGACAGCCCCGCGGTGAA-3

PHB2: 5’-CACCGGGCCCGAATGTCATAGATAACAAA-3’

### APEX Biotinylation assay

The APEX2 reaction was done under previously published conditions (Hung et al., 2014). Cells expressing APEX-KRAS construct were incubated in culture medium supplemented in 250μM Biotin-tyramide (A8011, APExBIO) for 30min at 37C°. 2 mM H_2_O_2_ was added and incubated for 1 min at room temperature. The reaction was quenched by three washes with PBS supplemented in 5mM Trolox (238813, Sigma Aldrich), 5mM of sodium ascorbate (PHR1279, Sigma Aldrich) and 1mM Sodium azide (S2002, Sigma Aldrich). A last wash was performed with PBS and the cells were immediately lysed on the dish with 150μL of RIPA (25 mM Tris-HCl pH 7.6, 150 mM NaCl, 1% NP-40, 1% sodium deoxycholate, 0.1% SDS) containing a EDTA-free protease inhibitor cocktail tablet (A32965, Thermoscientific) and phosphatase inhibitor tablet (A32957, Thermoscientifc). Cell lysates were dosed with Pierce BCA Protein Assay Kit and conserved at −80C°.

### Pull Down assay

For Pull down assays, 250 to 1000ng of proteins from cell lysates were incubated with 30 to 100 μL of RIPA-equilibrated Pierce Streptavidin magnetic beads (88816, ThermoScientific) at 4C° overnight. Supernatants were conserved at −80C°. For western blot analysis, beads were washed three times in RIPA. Elution was performed by adding Laemli buffer supplemented with 2mM Biotin and heated at 95C° for 5min.

For AP-MS assays, beads were washed twice with 1mL de RIPA, once with 1mL de KCL (1M), once with 1mL de Na2CO3 (0,1M), once with 1mL Urea (2M) diluted in Tris-HCl pH 8 (10mM) than once with TrisHCl pH7,5, NaCl 150mM, EDTA 1mM. Elution was performed by adding Laemli buffer supplemented with 2mM Biotin and heated at 95C° for 5min. 10% of the eluted sample were analyzed by western blotting for sample validation.

### Proteomics analysis

#### Sample preparation

Proteins were eluted from beads in 75 μL of Laemmli buffer under reducing and denaturing conditions. Bottom-up experiments’ tryptic peptides were obtained by Strap Micro Spin Column according to the manufacturer’s protocol (Protifi, NY, USA). Briefly: proteins from the above eluate were diluted 1:1 with 2x reducing-alkylating buffer (20 mM TCEP, 100 mM Chloroacetamide in 400 mM TEAB pH 8.5 and 4% SDS) and left 5 minutes at 95°C to allow reduction and alkylation in one step. Strap binding buffer was applied to precipitate proteins on quartz and proteolysis took place during 14h at 37°C with 1μg Trypsin sequencing grade (Promega). After speed-vacuum drying of eluted peptides, these were solubilized in 0.1% trifluoroacetic acid (TFA) in 10% Acetonitrile (ACN).

#### Liquid Chromatography-coupled Mass spectrometry analysis (LC-MS)

LC-MS analyses were performed on a Dionex U3000 HPLC nanoflow system coupled to a TIMS-TOF Pro mass spectrometer (Bruker Daltonik GmbH, Bremen, Germany). One μL was loaded, concentrated and washed for 3min on a C18 reverse phase precolumn (3μm particle size, 100 Å pore size, 75 μm inner diameter, 2 cm length, from Thermo Fisher Scientific). Peptides were separated on an Aurora C18 reverse phase resin (1.6 μm particle size, 100Å pore size, 75μm inner diameter, 25cm length mounted onto the Captive nanoSpray Ionisation module, (IonOpticks, Middle Camberwell Australia) with a 60 minutes overall run-time gradient ranging from 99% of solvent A containing 0.1% formic acid in milliQ-grade H2O to 40% of solvent B containing 80% acetonitrile, 0.085% formic acid in mQH2O. The mass spectrometer acquired data throughout the elution process and operated in DDA PASEF mode with a 1.1 second/cycle, with Timed Ion Mobility Spectrometry (TIMS) mode enabled and a data-dependent scheme with full MS scans in PASEF mode. This enabled a recurrent loop analysis of a maximum of the 120 most intense nLC-eluting peptides which were CID-fragmented between each full scan every 1.1 second. Ion accumulation and ramp time in the dual TIMS analyzer were set to 100 ms each and the ion mobility range was set from 1/K0 = 0.6 Vs cm-2 to 1.6 Vs cm-2. Precursor ions for MS/MS analysis were isolated in positive mode with the PASEF mode set to « on » in the 100-1.700 m/z range by synchronizing quadrupole switching events with the precursor elution profile from the TIMS device. The cycle duty time was set to 100%, accommodating as many MSMS in the PASEF frame as possible. Singly charged precursor ions were excluded from the TIMS stage by tuning the TIMS using the otof control software, (Bruker Daltonik GmbH). Precursors for MS/MS were picked from an intensity threshold of 2.500 arbitrary units (a.u.) and resequenced and summed until reaching a ‘target value’ of 20.000 a.u taking into account a dynamic exclusion of 0.40 s elution gap.

#### Protein quantification and comparison

The mass spectrometry data were analyzed using Maxquant version 1.6.17 (Tyanova et al., 2015). The database used was a concatenation of Homo sapiens sequences from the Swissprot databases (release june 2020: 563,972 sequences; 203,185,243 residues) and an in-house list of frequently found contaminant protein sequences. The enzyme specificity was trypsin’s. The precursor and fragment mass tolerances were set to 20ppm. Oxidation of methionines was set as variable modifications while carbamidomethylation of cysteines was considered complete. Second peptide search was allowed and minimal length of peptides was set at 7 amino acids. False discovery rate (FDR) was kept below 1% on both peptides and proteins. Label-free protein quantification (LFQ) was done using both unique and razor peptides. At least 2 such peptides were required for LFQ. For differential analysis, LFQ results from MaxQuant were quality-checked using PTXQC (Bielow et al., 2016). This work on MS was supported by the DIM Thérapie Génique Région Ile-de-France, IBiSA, and the Labex GR-Ex.

### Immunoblot analysis

Cells were resuspended in RIPA lysis buffer. For western blotting, following SDS–PAGE, proteins were transferred to 0.45 μM nitrocellulose membranes using Trans-BlotR Turbo™ Transfer System Cell system (Bio-Rad). The membrane was then blocked in 5% nonfat dry milk TBS 0.05% Tween 20 and incubated with primary antibody overnight at 4°C. Blots were incubated with the appropriate secondary antibodies for 1 h at room temperature and visualized using the Chemi-Doc XRS+ system (Bio-Rad).

### Immunocytochemistry

Cells were cultured on glass slides then fixed in PBS containing 4% paraformaldehyde and 4% sucrose for 15min. Cells were permeabilized for 5 min at room temperature in 0.25% Triton-X-100 in PBS, washed twice with PBS, and incubated for 30 min at 37°C in PBS containing 10% BSA. Cells were incubated overnight at 4°C with primary antibodies diluted in PBS containing 3% BSA. After washing, cells were incubated for 90 min at room temperature with the appropriate Alexa-conjugated secondary antibodies diluted in PBS containing 3% BSA. Cells were washed with PBS and mounted with ProLong Diamond Antifade Reagent with DAPI (Invitrogen, Carlsbad, CA, USA). Fluorescence images were acquired with Nikon A1 Rsi Inverted Confocal Microscope (Nikon, Tokyo, Japan) with NIS-Elements software (Nikon).

### Flow Cytometry assay

Cells were trypsinated then washed once in PBS. Cells were the fixed and permeabilized using BioLegend’s FOXP3 Fix/Perm Buffer (00-5523-00, Thermo Scientific) according to the manufacturer’s instructions. Cells were then sustained with recombinant APC coupled Anti-TOMM20 antibody (ab225341, Abcam) for 1h at room temperature. Celles were then washed once with perm-buffer and once with PBS. Flow cytometry analysis was performed just after staining.on an Accuri C6 flow cytometer (BD biosciences). Flow cytometry data were analysed using FlowJo software (BD Biosciences).

### Bioimaging Data analysis

Immunocytochemistry data were analyzed using an unbiased automatized program for colocalization analysis (SODA) (Lagache et al., 2018) and distance map analysis (Saito and Toriwaki, 1994) using ICY software (de Chaumont et al., 2012) (https://icy.bioimageanalysis.org). For Distance Map analysis, the distance between two channels is converted to a shade of gray image. Cell per cell intensity is then calculated. More the distance is low between those two channels, more the intensity is high. The sum of intensity per cell is then relativized by the area.

### Primary Antibodies

Recombinant Anti-Bcl-XL antibody – Monoclonal Rabbit – (ab31370) Abcam

Anti-Rab5 antibody - Early Endosome Marker - Polyclonal Rabbit - (ab18211) Abcam

Anti-PHB2 (E1Z5A) - Monoclonal Rabbit - (#14085) Cell Signaling

Recombinant Anti-TOMM20 antibody - Mitochondrial Marker – Monoclonal Mouse (ab186734) Abcam

mtHSP70 Antibody – Mouse Monoclonal - (MA3-028) Invitrogen

p44/42 MAPK (Erk1/2) Antibody – Rabbit Polyclonal - (#9102) Cell Signaling

Phospho-p44/42 MAPK (Erk1/2) (Thr202/Tyr204) (197G2) - Rabbit Monoclonal-(#4377) Cell Signaling

anti-Actin antibody – Mouse Monoclonal - (MAB1501) Millipore

Anti-β-Tubulin antibody - Mouse monoclonal - Sigma-Aldrich

### Secondary Antibodies

Horseradish Peroxidase AffiniPure Goat Anti-Mouse IgG (H+L) – Jackson IR

Horseradish Peroxidase AffiniPure Goat Anti-Rabbit IgG (H+L) – Jackson IR

Goat anti-Rabbit IgG (H+L) Highly Cross-Adsorbed Secondary Antibody, Alexa Fluor™ 568 (A-11036, Thermo Scientific)

Goat anti-Mouse IgG (H+L) Highly Cross-Adsorbed Secondary Antibody, Alexa Fluor™ Plus 647 (A32728, Thermo Scientific)

### Fluorescent probes

Streptavidin, Alexa Fluor™ 647 conjugate (S21374, Thermo Scientific)

MitoTracker™ Deep Red FM (M22426, Thermo Scientific)

## Acknowledgements

NB was supported by a fellowship from Ligue contre le Cancer. This work was supported by INCa and DGOS (SIRIC ILIAD, INCa-DGOS Inserm-12558) and Ligue contre le cancer (CD44, CD53 and CD22)

We thank the IBISA MicroPICell facility (SFR Bonamy, Biogenouest), member of the national infrastructure France-Bioimaging supported by the French national research agency (ANR-10-INBS-04) and the Cytocell Cytometry Facility (SFR Bonamy) from Nantes for invaluable advice and expert technical assistance.

---

**Supplementary Figure 1.**
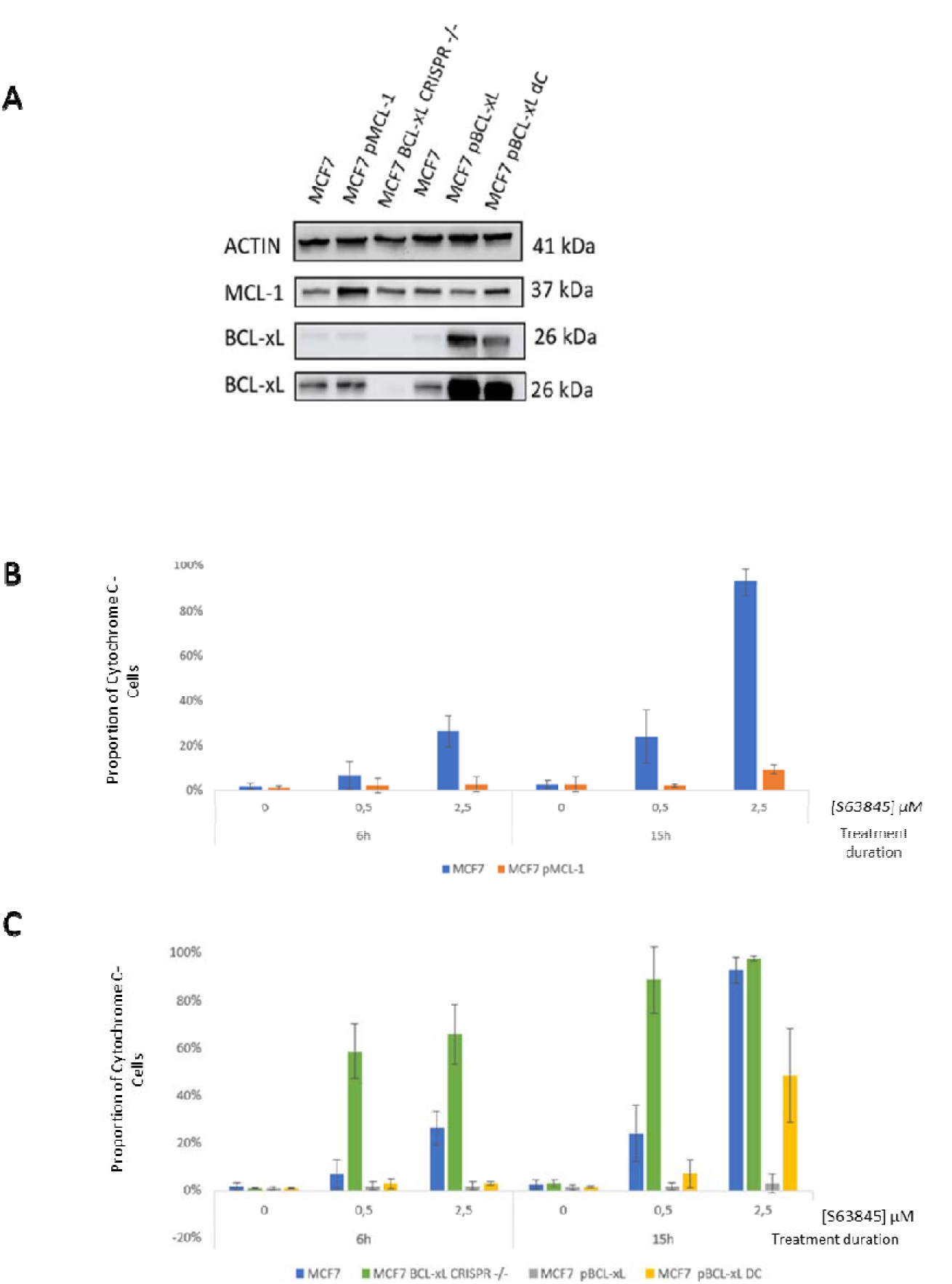
**A** Western blot analysis of BCL-xL and MCL-1 expression in MCF7, MCF7 pLVX-MCL-1, MCF7 BCL-xL CRISPR, MCF7 pLVX-BCL-xL or MCF7pLVX-BCL-xL ΔC (C-terminal depleted BCL-xL, or BCL-xLdC) cell lines **B** MOMP estimated by the proportion of cells showing Cytochrome C release measured by Flux Cytometry in the MCF7 and MCF7 pLVX-MCL-1. **C**. MOMP estimated by the proportion of cells harbouring Cytochrome C release measured by Flux Cytometry in the MCF7, MCF7 BCL-xL CRISPR, MCF7pLVX-BCL-xL or MCF7pLVX-BCL-xL ΔC cell lines.

**Supplementary Figure 2.**
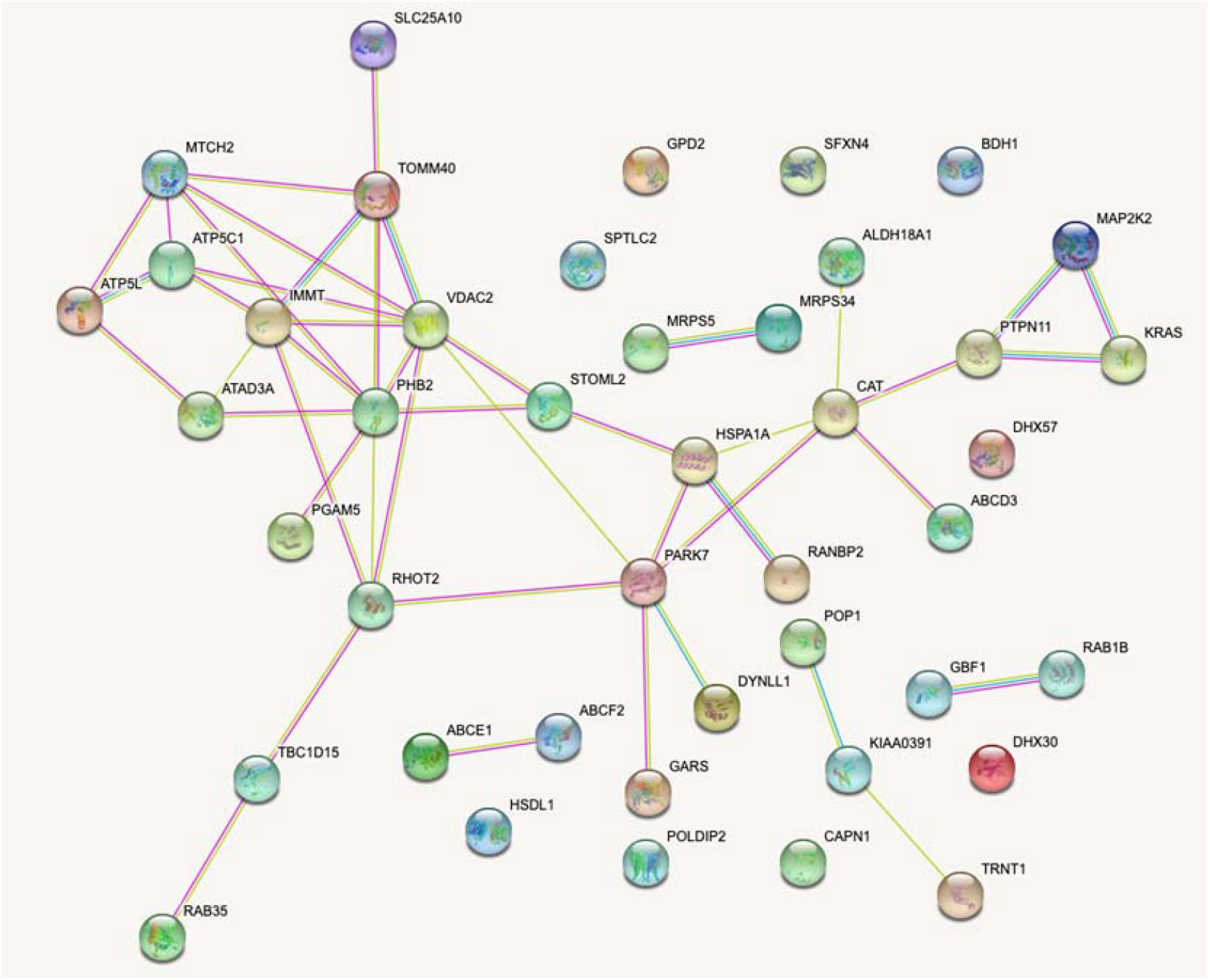
String Database analysis of the 43 mitochondrial APEX-KRAS proximitome in MCF7 BCL-xL CRISPR after Mass Spectrometry detection and Fold Change analysis

## References

Amendola, C.R., Mahaffey, J.P., Parker, S.J., Ahearn, I.M., Chen, W.-C., Zhou, M., Court, H., Shi, J., Mendoza, S.L., Morten, M., et al. (2019). KRAS4A Directly Regulates Hexokinase 1. Nature 576, 482–486. https://doi.org/10.1038/s41586-019-1832-9.

Bertolin, G., Alves-Guerra, M.-C., Cheron, A., Burel, A., Prigent, C., Le Borgne, R., and Tramier, M. (2021). Mitochondrial Aurora kinase A induces mitophagy by interacting with MAP1LC3 and Prohibitin 2. Life Sci. Alliance 4, e202000806. https://doi.org/10.26508/lsa.202000806.

Bielow, C., Mastrobuoni, G., and Kempa, S. (2016). Proteomics Quality Control: Quality Control Software for MaxQuant Results. J. Proteome Res. 15, 777–787. https://doi.org/10.1021/acs.jproteome.5b00780.

Bivona, T.G., Quatela, S.E., Bodemann, B.O., Ahearn, I.M., Soskis, M.J., Mor, A., Miura, J., Wiener, H.H., Wright, L., Saba, S.G., et al. (2006). PKC regulates a farnesyl-electrostatic switch on K-Ras that promotes its association with Bcl-XL on mitochondria and induces apoptosis. Mol. Cell 21, 481–493. https://doi.org/10.1016/j.molcel.2006.01.012.

Braun, F., de Carné Trécesson, S., Bertin-Ciftci, J., and Juin, P. (2013). Protect and serve: Bcl-2 proteins as guardians and rulers of cancer cell survival. Cell Cycle 12, 2937–2947. https://doi.org/10.4161/cc.25972.

Carné Trécesson, S. de, Souazé, F., Basseville, A., Bernard, A.-C., Pécot, J., Lopez, J., Bessou, M., Sarosiek, K.A., Letai, A., Barillé-Nion, S., et al. (2017). BCL-XL directly modulates RAS signalling to favour cancer cell stemness. Nat. Commun. 8, 1123. https://doi.org/10.1038/s41467-017-01079-1.

de Chaumont, F., Dallongeville, S., Chenouard, N., Hervé, N., Pop, S., Provoost, T., Meas-Yedid, V., Pankajakshan, P., Lecomte, T., Le Montagner, Y., et al. (2012). Icy: an open bioimage informatics platform for extended reproducible research. Nat. Methods 9, 690–696. https://doi.org/10.1038/nmeth.2075.

Chen, X., Glytsou, C., Zhou, H., Narang, S., Reyna, D.E., Lopez, A., Sakellaropoulos, T., Gong, Y., Kloetgen, A., Yap, Y.S., et al. (2019). Targeting Mitochondrial Structure Sensitizes Acute Myeloid Leukemia to Venetoclax Treatment. Cancer Discov. 9, 890–909. https://doi.org/10.1158/2159-8290.CD-19-0117.

Cho, K., Park, J.-H., Piggott, A.M., Salim, A.A., Gorfe, A.A., Parton, R.G., Capon, R.J., Lacey, E., and Hancock, J.F. (2012). Staurosporines disrupt phosphatidylserine trafficking and mislocalize Ras proteins. J. Biol. Chem. 287, 43573–43584. https://doi.org/10.1074/jbc.M112.424457.

Hammerling, B.C., Najor, R.H., Cortez, M.Q., Shires, S.E., Leon, L.J., Gonzalez, E.R., Boassa, D., Phan, S., Thor, A., Jimenez, R.E., et al. (2017). A Rab5 endosomal pathway mediates Parkin-dependent mitochondrial clearance. Nat. Commun. 8, 14050. https://doi.org/10.1038/ncomms14050.

Hernando-Rodríguez, B., and Artal-Sanz, M. (2018). Mitochondrial Quality Control Mechanisms and the PHB (Prohibitin) Complex. Cells 7, E238. https://doi.org/10.3390/cells7120238.

Hung, V., Zou, P., Rhee, H.-W., Udeshi, N.D., Cracan, V., Svinkina, T., Carr, S.A., Mootha, V.K., and Ting, A.Y. (2014). Proteomic mapping of the human mitochondrial intermembrane space in live cells via ratiometric APEX tagging. Mol. Cell 55, 332–341. https://doi.org/10.1016/j.molcel.2014.06.003.

Juin, P., Geneste, O., Gautier, F., Depil, S., and Campone, M. (2013). Decoding and unlocking the BCL-2 dependency of cancer cells. Nat. Rev. Cancer 13, 455–465. https://doi.org/10.1038/nrc3538.

Lagache, T., Grassart, A., Dallongeville, S., Faklaris, O., Sauvonnet, N., Dufour, A., Danglot, L., and Olivo-Marin, J.-C. (2018). Mapping molecular assemblies with fluorescence microscopy and object-based spatial statistics. Nat. Commun. 9, 698. https://doi.org/10.1038/s41467-018-03053-x.

Lee, D.-H., Kim, D., Kim, S.T., Jeong, S., Kim, J.L., Shim, S.M., Heo, A.J., Song, X., Guo, Z.S., Bartlett, D.L., et al. (2018). PARK7 modulates autophagic proteolysis through binding to the N-terminally arginylated form of the molecular chaperone HSPA5. Autophagy 14, 1870–1885. https://doi.org/10.1080/15548627.2018.1491212.

Lee, S.-Y., Kang, M.-G., Park, J.-S., Lee, G., Ting, A.Y., and Rhee, H.-W. (2016). APEX Fingerprinting Reveals the Subcellular Localization of Proteins of Interest. Cell Rep. 15, 1837–1847. https://doi.org/10.1016/j.celrep.2016.04.064.

Liu, J., Zhang, C., Zhao, Y., Yue, X., Wu, H., Huang, S., Chen, J., Tomsky, K., Xie, H., Khella, C.A., et al. (2017). Parkin targets HIF-1α for ubiquitination and degradation to inhibit breast tumor progression. Nat. Commun. 8, 1823. https://doi.org/10.1038/s41467-017-01947-w.

Lu, A., Tebar, F., Alvarez-Moya, B., López-Alcalá, C., Calvo, M., Enrich, C., Agell, N., Nakamura, T., Matsuda, M., and Bachs, O. (2009). A clathrin-dependent pathway leads to KRas signaling on late endosomes en route to lysosomes. J. Cell Biol. 184, 863–879. https://doi.org/10.1083/jcb.200807186.

Mellacheruvu, D., Wright, Z., Couzens, A.L., Lambert, J.-P., St-Denis, N., Li, T., Miteva, Y.V., Hauri, S., Sardiu, M.E., Low, T.Y., et al. (2013). The CRAPome: a Contaminant Repository for Affinity Purification Mass Spectrometry Data. Nat. Methods 10, 730–736. https://doi.org/10.1038/nmeth.2557.

Morgenstern, M., Stiller, S.B., Lübbert, P., Peikert, C.D., Dannenmaier, S., Drepper, F., Weill, U., Höß, P., Feuerstein, R., Gebert, M., et al. (2017). Definition of a High-Confidence Mitochondrial Proteome at Quantitative Scale. Cell Rep. 19, 2836–2852. https://doi.org/10.1016/j.celrep.2017.06.014.

Polier, G., Neumann, J., Thuaud, F., Ribeiro, N., Gelhaus, C., Schmidt, H., Giaisi, M., Köhler, R., Müller, W.W., Proksch, P., et al. (2012). The Natural Anticancer Compounds Rocaglamides Inhibit the Raf-MEK-ERK Pathway by Targeting Prohibitin 1 and 2. Chem. Biol. 19, 1093–1104. https://doi.org/10.1016/j.chembiol.2012.07.012.

Priault, M., Hue, E., Marhuenda, F., Pilet, P., Oliver, L., and Vallette, F.M. (2010). Differential dependence on Beclin 1 for the regulation of pro-survival autophagy by Bcl-2 and Bcl-xL in HCT116 colorectal cancer cells. PloS One 5, e8755. https://doi.org/10.1371/journal.pone.0008755.

Rajalingam, K., Wunder, C., Brinkmann, V., Churin, Y., Hekman, M., Sievers, C., Rapp, U.R., and Rudel, T. (2005). Prohibitin is required for Ras-induced Raf-MEK-ERK activation and epithelial cell migration. Nat. Cell Biol. 7, 837–843. https://doi.org/10.1038/ncb1283.

Saito, T., and Toriwaki, J.-I. (1994). New algorithms for euclidean distance transformation of an n-dimensional digitized picture with applications. Pattern Recognit. 27, 1551–1565. https://doi.org/10.1016/0031-3203(94)90133-3.

Sung, P.J., Tsai, F.D., Vais, H., Court, H., Yang, J., Fehrenbacher, N., Foskett, J.K., and Philips, M.R. (2013). Phosphorylated K-Ras limits cell survival by blocking Bcl-xL sensitization of inositol trisphosphate receptors. Proc. Natl. Acad. Sci. U. S. A. 110, 20593–20598. https://doi.org/10.1073/pnas.1306431110.

Surve, S.V., Myers, P.J., Clayton, S.A., Watkins, S.C., Lazzara, M.J., and Sorkin, A. (2019). Localization dynamics of endogenous fluorescently labeled RAF1 in EGF-stimulated cells. Mol. Biol. Cell 30, 506–523. https://doi.org/10.1091/mbc.E18-08-0512.

Tyanova, S., Temu, T., Carlson, A., Sinitcyn, P., Mann, M., and Cox, J. (2015). Visualization of LC-MS/MS proteomics data in MaxQuant. Proteomics 15, 1453–1456. https://doi.org/10.1002/pmic.201400449.

Wei, G., Margolin, A.A., Haery, L., Brown, E., Cucolo, L., Julian, B., Shehata, S., Kung, A.L., Beroukhim, R., and Golub, T.R. (2012). Chemical Genomics Identifies Small-Molecule MCL1 Repressors and BCL-xL as a Predictor of MCL1 Dependency. Cancer Cell 21, 547–562. https://doi.org/10.1016/j.ccr.2012.02.028.

Yan, C., Gong, L., Chen, L., Xu, M., Abou-Hamdan, H., Tang, M., Désaubry, L., and Song, Z. (2020). PHB2 (prohibitin 2) promotes PINK1-PRKN/Parkin-dependent mitophagy by the PARL-PGAM5-PINK1 axis. Autophagy 16, 419–434. https://doi.org/10.1080/15548627.2019.1628520.

Yoshinaka, T., Kosako, H., Yoshizumi, T., Furukawa, R., Hirano, Y., Kuge, O., Tamada, T., and Koshiba, T. (2019). Structural Basis of Mitochondrial Scaffolds by Prohibitin Complexes: Insight into a Role of the Coiled-Coil Region. IScience 19, 1065–1078. https://doi.org/10.1016/j.isci.2019.08.056.

Yurugi, H., Marini, F., Weber, C., David, K., Zhao, Q., Binder, H., Désaubry, L., and Rajalingam, K. (2017). Targeting prohibitins with chemical ligands inhibits KRAS-mediated lung tumours. Oncogene 36, 4778–4789. https://doi.org/10.1038/onc.2017.93.

